# Generalizing Stepping Concepts To Non-Straight Walking

**DOI:** 10.1101/2023.05.15.540644

**Authors:** Jonathan B. Dingwell, Anna C. Render, David M. Desmet, Joseph P. Cusumano

## Abstract

People rarely walk in straight lines. Instead, we make frequent turns or other maneuvers. Spatiotemporal parameters fundamentally characterize gait. For straight walking, these parameters are well-defined for that task of walking on a straight *path*. Generalizing these concepts to *non*-straight walking, however, is not straightforward. People also follow non-straight paths imposed by their environment (store aisle, sidewalk, etc.) or choose readily-predictable, stereotypical paths of their own. People actively maintain lateral position to stay on their path and readily adapt their stepping when their path changes. We therefore propose a conceptually coherent convention that defines step lengths and widths relative to known walking paths. Our convention simply re-aligns lab-based coordinates to be tangent to a walker’s path at the mid-point between the two footsteps that define each step. We hypothesized this would yield results both more *correct* and more consistent with notions from straight walking. We defined several common non-straight walking tasks: single turns, lateral lane changes, walking on circular paths, and walking on arbitrary curvilinear paths. For each, we simulated idealized step sequences denoting “perfect” performance with known constant step lengths and widths. We compared results to path-independent alternatives. For each, we directly quantified accuracy relative to known *true* values. Results strongly confirmed our hypothesis. Our convention returned vastly smaller errors and introduced *no* artificial stepping asymmetries across all tasks. All results for our convention rationally generalized concepts from straight walking. Taking walking paths explicitly into account as important task goals themselves thus resolves conceptual ambiguities of prior approaches.

## 1. Introduction

When we walk in the real world, we rarely walk in straight lines. We frequently turn and/or make other maneuvers (Glaister et al., 2007), starting from when we first learn to walk (Adolph et al., 2012). As adults, we skillfully modulate foot placements (Bruijn and van Dieën, 2018) to maneuver (Acasio et al., 2017; Desmet et al., 2022). But as we get older, we are less able to shift our weight effectively (Ambrose et al., 2013), which can induce falls and serious injury (Burns and Kakara, 2018). Older adults fall most often while walking (Kelsey et al., 2012) and frequently while attempting non-straight walking tasks or maneuvers (Robinovitch et al., 2013).

Spatiotemporal gait parameters (step length, width, time, etc.) fundamentally characterize gait. People modify them regularly in real environments (Twardzik et al., 2019) and they can predict consequences like fall risk (Taylor et al., 2013). Spatiotemporal parameters are well-defined for straight walking (Richards et al., 2023), where most gait labs presume a constant forward direction of progression (DoP). Here, step lengths and widths are rightly defined relative to the task of walking on that straight path. However, generalizing these concepts to *non*-straight walking is not straightforward. Given the need to predict and/or prevent falls that often occur during complex tasks (Robinovitch et al., 2013) and increasing emphasis on real-world walking (Hafer et al., 2023), we must reassess what these biomechanical quantities mean for non-straight walking.

Some have proposed defining DoP from anatomically-based local coordinates, like those defined by center-of-mass (CoM) trajectories (Ho et al., 2023), or horizontal pelvis (Ho et al., 2023; Kainz et al., 2016) or trunk (Jansen et al., 2011) rotations. Unfortunately, these are decidedly problematic. Even for straight walking, the CoM continuously oscillates side-to-side (Saunders et al., 1953; Tesio et al., 2010), the pelvis continuously rotates (Lewis et al., 2017; Saunders et al., 1953), and the trunk rotates separately from the pelvis (Prins et al., 2019).

Thus, these approaches necessarily yield *many* possible (but arbitrary) DoPs that only rarely align with the true DoP (see Supplement). Such difficulties make clear this is not about merely finding the right “method”, but a fundamental question of how we conceive of these quantities beforehand.

An alternative based only on step placements was proposed by Huxham et al. (2006), extending Courtine & Schieppati (2003). Huxham calculates step lengths and widths from a DoP defined by a line connecting consecutive ipsilateral foot placements. This convention has been applied to executing turns in one step (He et al., 2018; Huxham et al., 2006) or over multiple steps (Conradsson et al., 2018; Courtine and Schieppati, 2003; Dixon et al., 2013; Tillman et al., 2022), to walking around circular (Ho et al., 2023; Xu et al., 2017) or other non-straight (Bland et al., 2019; Fino et al., 2016) paths, even walking outdoors (Bergsma et al., 2021). However, Huxham’s convention produces several counter-intuitive consequences. First, consecutive step lengths are measured along different DoP (see their Fig. 1B). Hence, adding consecutive step lengths will not equal the corresponding stride length, inconsistent with definitions from straight walking. Second, Huxham’s convention exaggerates distinctions between so-called “step turn” steps (where the stepping foot steps away from the stance foot) and “spin turn” steps (where the stepping foot steps towards/across the stance foot) (Hase and Stein, 1999). While seemingly relevant for single-step turns, when people turn over multiple steps, they necessarily alternate “step turn” steps with “spin turn” steps, making the idea of differentiating these as distinct “strategies” (Conradsson et al., 2018; Dixon et al., 2013) somewhat illogical. Even for straight walking, people step slightly away from or towards/across their stance foot at every step, again making Huxham’s convention inconsistent with conventional definitions. Third, not all non-straight walking involves turns, as sometimes we step sideways to make lateral maneuvers (Acasio et al., 2017; Desmet et al., 2022). It is not clear if Huxham’s convention can even be meaningfully applied to such tasks.

**Figure 1.**
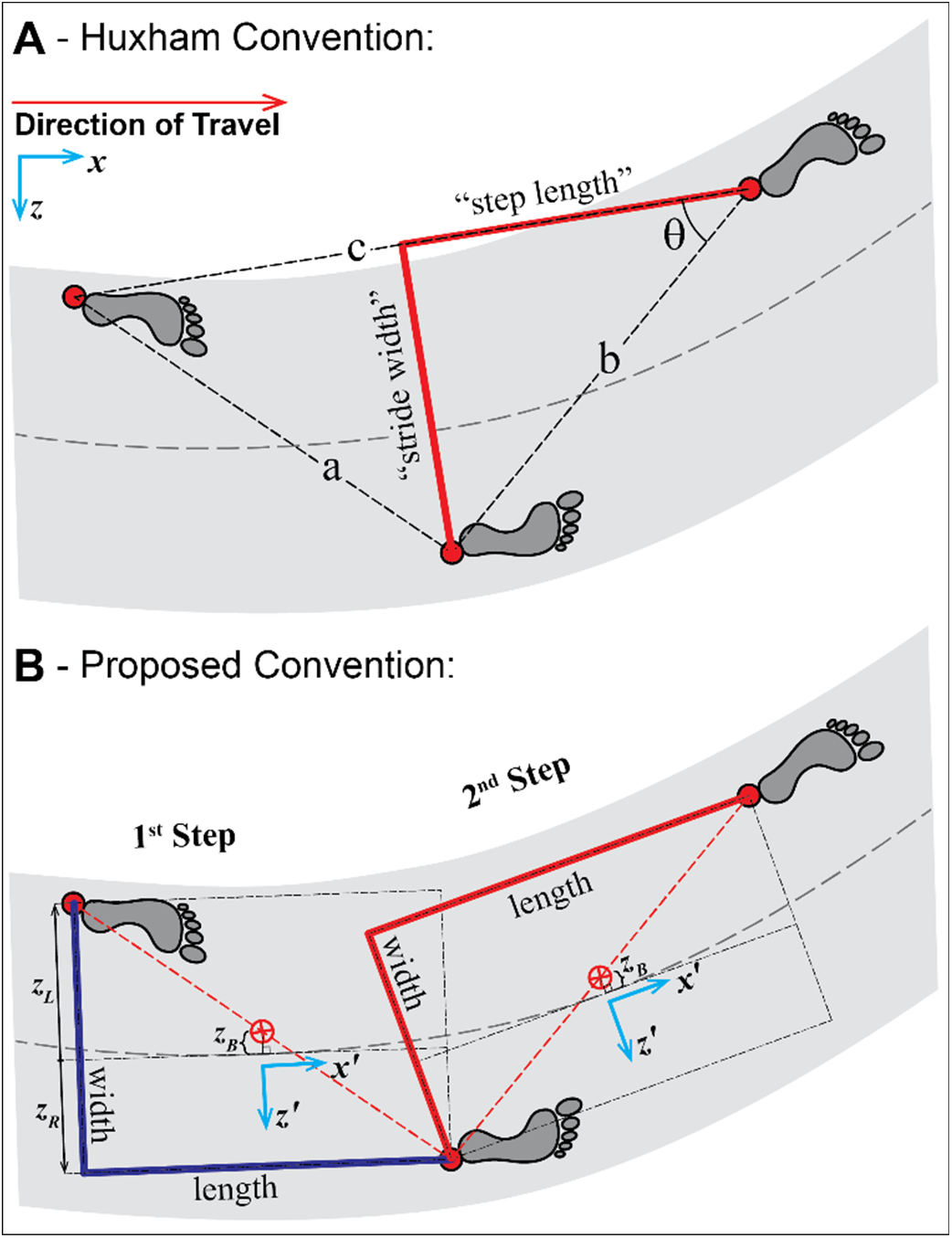
Defining Stepping Parameters: **A)** The convention for defining step length and “stride width” proposed by Huxham et al., 2006. Given only the locations of three consecutive foot placements, the direction of forward progression (DoP) for the middle foot placement is taken as line “*c*”, which connects the prior and subsequent contralateral foot placements. Huxham’s step lengths (*l*_*H*_) and “stride widths” (*w*_*H*_) are then computed from basic trigonometric relationships: i.e., *l*_*H*_ = *b*cos(θ) = (*b*^2^+*c*^2^−*a*^2^)/2*c* and *w*_*H*_ = sqrt(*b*^2^−*l*_*H*_^2^) (Huxham et al, 2006). Of note, these relationships do not in any way account for where the feet are placed with respect to the walking path. **B)** The proposed convention presenting a path-based approach for the exact same three foot placements. Each step is considered its own true step, consisting of two foot placements only. The person’s body position is taken as the midpoint between the two foot placements that make up that step (Orendurff et al., 2004; Dingwell & Cusumano, 2019). A local [*x*′,*z*′] coordinate system is then aligned tangent to the path at the point on the path closest to the body position. Step lengths (*l*_*P*_) and step widths (*w*_*P*_) are then calculated in this path-defined local coordinate system just as they would be in fixed lab-based coordinates for typical straight walking.

It is important to view stepping (and hence, walking) as not merely something to be “measured”, but the product of a *task* one performs. The prior efforts above, however, did not consider that people walk intentionally (Gordon et al., 2021) to achieve specific goal(s) (Cusumano and Cesari, 2006). People follow *paths* (store aisles, sidewalks, etc.) when they walk (Dingwell and Cusumano, 2019). When their environment does not impose an explicit path, people choose their own (Moussaïd et al., 2011). Indeed, these paths are stereotypical (Hicheur et al., 2007) and well-predicted by conventional control laws (Arechavaleta et al., 2008; Brown et al., 2021). People then balance maintaining step width with their lateral position relative to their path, so they do not drift sideways off their path (Dingwell and Cusumano, 2019). They readily modulate how they regulate their steps if given explicit feedback (Render et al., 2021), or to make lateral maneuvers (Desmet et al., 2022). Conversely, neither Huxham’s convention nor the anatomically-based approaches described above reference the path followed or any other goal(s) to maintain. The logic of those approaches thus seems backwards: tasks are not created by where people step – instead, people choose where to step to accomplish particular task goals (including to follow some path). Because stepping behavior is well-explained by regulation processes that minimize errors with respect to specific task goals (Desmet et al., 2022; Dingwell and Cusumano, 2019; Dingwell et al., 2010), stepping parameters (step length, width, etc.) should be defined consistently with these processes.

Referencing stepping parameters to the walking path/trajectory directly generalizes original stepping notions from straight walking (Richards et al., 2023) and also respects the nature of intrinsic motor regulation. As such, we hypothesize this will yield more *accurate* results. We first propose a convention for defining step lengths and widths relative to a known path. We compare our proposed approach to that of Huxham et al. (2006). Any measure’s accuracy can only be defined relative to known true values. As true “correct” values can never be known for experimental data, we instead simulated idealized step sequences to constitute “perfect” performance in several non-straight walking tasks. We show our proposed approach fully disambiguates “steps” from “strides”, yields more accurate measures that are far more consistent with straight walking definitions, applies across multiple non-straight tasks, and accommodates primary stepping goals people try to maintain as they walk.

## 2. Methods

### 2.1 Stepping Parameters for Non-Straight Walking

The challenge is to define step lengths and widths relative to some direction of forward progression (DoP) that may change at each step (Huxham et al., 2006). The convention proposed by Huxham et al. (2006) defines a DoP for each *stride* as the line connecting the two ipsilateral heel strikes that form that stride (Fig. 1A). “Stride width” (*w*_*H*_) is measured perpendicular to this line. Step length (*l*_*H*_) is measured from the resulting intersection to the heel of the leading foot (Fig. 1A). These measurements make no reference to any path the person walks on or follows.

However, people do not “just walk” – they walk intentionally (Gordon et al., 2021) in goal-directed ways (Cusumano and Cesari, 2006). As such, they follow paths to reach their destinations (Arechavaleta et al., 2008; Moussaïd et al., 2011). Given any such path, here we define each individual step from only its two consecutive foot placements (Fig. 1B). We take body position for that step as the mid-point between them (Dingwell and Cusumano, 2019). This can serve as a discrete (once per step) proxy for CoM location (Desmet et al., 2022; Orendurff et al., 2004), but has the virtue of being determined entirely from step positions. Then, just as one would transform local coordinate systems between body segments (foot-to-shank, shank-to-thigh, etc.), we merely rotate the global coordinate system, [*x,z*], to be tangent to the path at the point nearest to the body position ([*x′,z′*]; Fig. 1B). We then calculate step widths (*w*_*P*_) and lengths (*l*_*P*_) as usual (Richards et al., 2023), only now in path-aligned [*x′,z′*] coordinates (Fig. 1B). We define foot placements ([*x′*_*L*_,*z′*_*L*_] and [*x′*_*R*_,*z′*_*R*_]) relative to the relevant tangent point on the path. Step length is the distance in +*x′* from trailing to leading foot. Step width is the lateral displacement between the right and left feet: *w* = *z′*_*R*_ − *z′*_*L*_ (i.e., so cross-over steps yield negative *w*). We additionally define lateral body position relative to the path (Dingwell and Cusumano, 2019) (not analyzed here) as the locally-lateral midpoint between the feet: *z′*_*B*_ = 1/2(*z′*_*L*_ + *z′*_*R*_).

### 2.2 Non-Straight Walking Tasks

We defined several non-straight walking tasks humans commonly perform: make a *Single Turn* (as though turning a corner), make a *Lateral Lane Change* (such as to avoid an obstacle), walk around a *Circular Path* (as though going around a curve), and walk along an *Arbitrary Curvilinear Path* (such as a winding garden path). For all tasks, we assumed a global (i.e., lab) coordinate system as [*x,z*] in the horizontal plane to be +*x* “forward” (left-to-right in figures) and +*z* to the walker’s right (down in figures).

For each task, we simulated idealized step sequences that constituted “perfect” performance (i.e., zero error) in that task. We constructed step sequences with all steps having the same pre-defined step length (*L* = 0.60m) and step width (*W* = 0.15m). While these values are typical (Herssens et al., 2020), our findings and conclusions do not depend on any specific choice(s) of *L* and/or *W*.

For *Single Turns*, we defined a path to run straight along the +*x*-axis, then (instantaneously) turn by angle Θ_Turn_ at the origin ([0,0]), then continue straight again in the new direction. We derived, as geometric functions of *L, W*, and Θ_Turn_, where to place each foot so every step had length *L* and width *W* and was located exactly ±*W*/2 to either side of the path. Hence, the walker walks with exactly *L* and *W* up to the turn step, turns the full Θ_Turn_ at that foot placement, then continues walking with exactly *L* and *W* in the new (turned) direction. We present results here for Θ_Turn_ ∈ {−30°, −75°, +60°}.

For *Lateral Lane Change* maneuvers, we defined a path to run straight along the +*x*-axis, with a lateral shift of displacement Δ*z* occurring just after the origin ([0,0]), and then continuing straight in the +*x* direction after the lane-change. We derived, as functions of *L, W*, and Δ*z*, where to place each foot so every step had length *L* and width *W* and was exactly ±*W*/2 to either side of the path. Here, the walker walks with steps of exactly *L* and *W* up to the origin, fully changes lanes on the next foot placement, then continues walking with exactly *L* and *W* thereafter. We present results here for Δ*z* ∈ {+30cm, +60cm, −45cm} (or in terms of step widths: Δ*z* ∈ {+2, +4,−3}·*W*).

For walking around *Circular Paths*, we defined each path as a circle centered at the origin with radius *R*_*Path*_. We placed each alternating footstep along the path at consecutive distances *L*, measured along the arc of the circle, and with alternating feet placed at radial distances *R*_*Path*_±*W*/2 perpendicular to the circular path. Here, we took these footstep sequences to denote walking counter-clockwise around each circle. However, the exact same footsteps could equally represent clockwise walking. We present results here for individual paths of *R*_*Path*_ ∈{6.0m, 1.5m, 0.75m} and also across the range of 0.5m ≤ *R*_*Path*_ ≤ 10.0m.

For walking along an *Arbitrary Curvilinear Path*, we defined a path as a 5^th^ order polynomial:

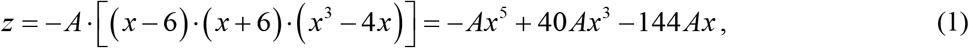

on the interval *x* ∈ [−6m, +6m] and with amplitude *A* = 0.0025m. This yielded a path that was symmetrical about the origin, started and ended on the +*x*-axis, and had two larger peaks (minimum radius-of-curvature (i.e., instantaneous *R*_*Path*_) ≈ 0.61m) and two smaller peaks (minimum radius-of-curvature ≈ 1.6m). The shape and curvature range of this path are typical of real paths (Arechavaleta et al., 2008; Rudenko et al., 2020). We placed each alternating footstep at consecutive distances *L* measured along the polynomial, and with alternating feet placed at distances ±*W*/2 perpendicular to the path at each foot placement. This yielded a sequence of 32 consecutive footsteps along this path.

### 2.3 Data Analyses

For all steps in each simulated task, we used Huxham’s convention (Fig. 1A) to compute step lengths (*l*_*H*_) and stride widths (*w*_*H*_). We used our proposed convention (Fig. 1B) to compute step lengths (*l*_*P*_) and widths (*w*_*P*_). To quantify the *accuracy* of each measurement, we scaled each computed value to the pre-defined *true* values, *L* and *W*: any value {*l*_*H*_,*l*_*P*_} ≠ *L* or {*w*_*H*_,*w*_*P*_} ≠ *W* indicated an error. We also calculated percent errors (%Error) in each step measure as:

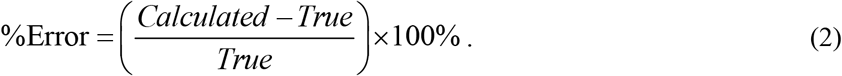

For circular paths, we computed and plotted %Errors of all measures across the range of 0.5m ≤ *R*_*Path*_ ≤ 10.0m for both “outside-to-inside” and “inside-to-outside” steps. We computed a Universal Symmetry Index (*USI*) (Alves et al., 2020) between these outside-to-inside (“*χ*_*oi*_”) and inside-to-outside (“*χ*_*io*_”) steps as:

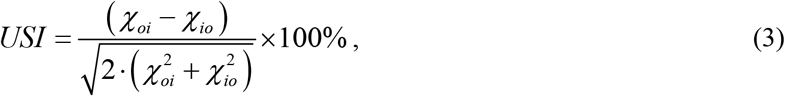

where *χ* ∈ {*l*_*H*_, *l*_*P*_, *w*_*H*_, *w*_*P*_}. For {*χ*_*oi*_, *χ*_*io*_} that can have either positive or negative values, Eq. (3) returns *USI* = 0 for perfect symmetry (*χ*_*oi*_ = *χ*_*io*_), with a bounded maximal range of *USI* ∈ [−100%,+100%] otherwise (Alves et al., 2020). Because we simulated perfectly aligned steps along each circular path, step lengths and widths should likewise be perfectly symmetrical: i.e., one should obtain the same results for all steps, whether walking clockwise or counter-clockwise around the circle.

Unlike for straight paths, setting [*x′,z′*] at different locations along any *curved* path will alter resulting step widths and step lengths calculated, potentially introducing (artificial) asymmetries (Fig. 2). Here, we directly quantified these effects for our proposed convention. We computed *l*_*P*_ and *w*_*P*_ across the range of possible [*x′,z′*] from the trailing (rear) foot placement (Fig. 2A) to the leading (front) foot placement (Fig. 2B) for circular paths of *R*_*Path*_ ∈{6.0m, 1.5m, 0.75m}. At each [*x′,z′*], we computed %Errors in both step length and width for both outside-to-inside and inside-to-outside steps and *USI* values (Eq. 3) between them.

**Figure 2.**
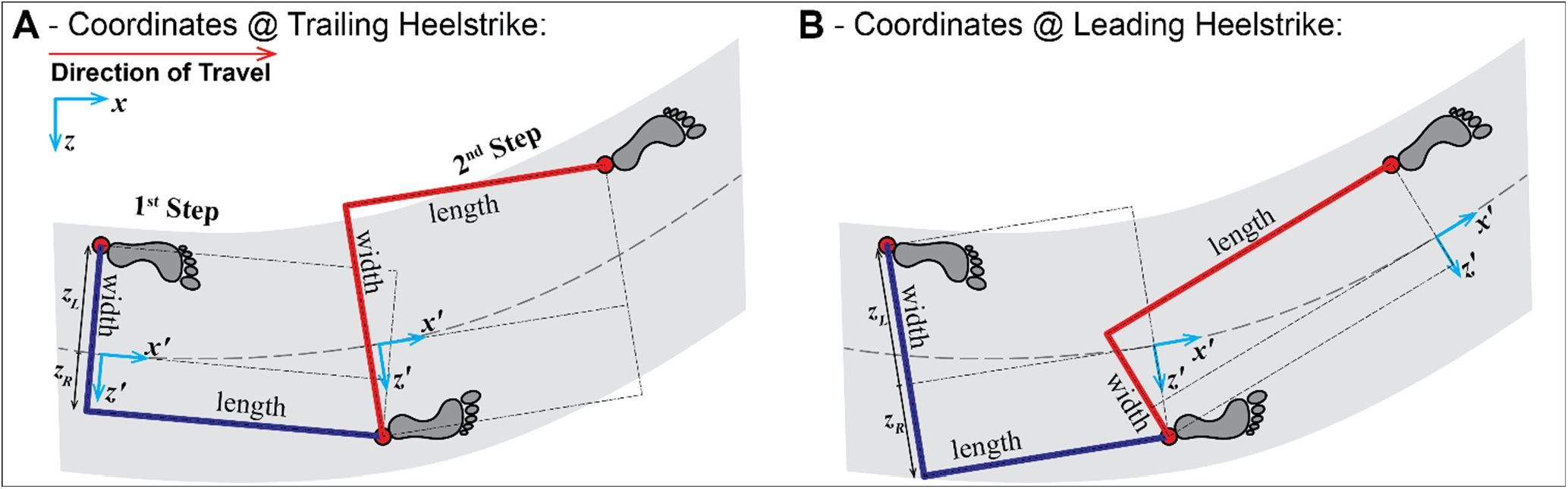
Different Local Path Coordinate Choices. When [*x′,y′*] are aligned with the midpoint between the two feet (Fig. 1B), step lengths and step widths are each similar for both steps. However, **A)** If [*x′,y′*] were instead aligned with the trailing (rear) foot placement of each step, the 1^st^ (“inside-to-outside”) step becomes longer and narrower, while the 2^nd^ (“outside-to-inside”) step becomes wider and shorter. **B)** Conversely, if [*x′,y′*] were aligned with the leading (front) foot placement of each step, the 1^st^ step now becomes wider and shorter, while the 2^nd^ step becomes longer and narrower.

For the arbitrary curvilinear path, we computed %Errors for both measures at each step and plotted these versus the *x* axis location of each corresponding step.

## 3. Results

For *Single Turns* (Fig. 3), both conventions yielded perfect performance ({*l*_*H*_,*l*_*P*_} = *L*; {*w*_*H*_,*w*_*P*_} = *W*) for the straight sections of all paths (not marked on figure). For the turn step itself, however, Huxham’s convention (Fig. 3A) yielded substantial errors in step length (*l*_*H*_: −36% ≤ %Errors ≤ −1%) and especially step width (*w*_*H*_: −213% ≤%Errors ≤ +223%). Conversely, our convention (Fig. 3B) yielded *no* errors at any step for any turn angle (*l*_*P*_ &*w*_*P*_: all %Error = 0%).

**Figure 3.**
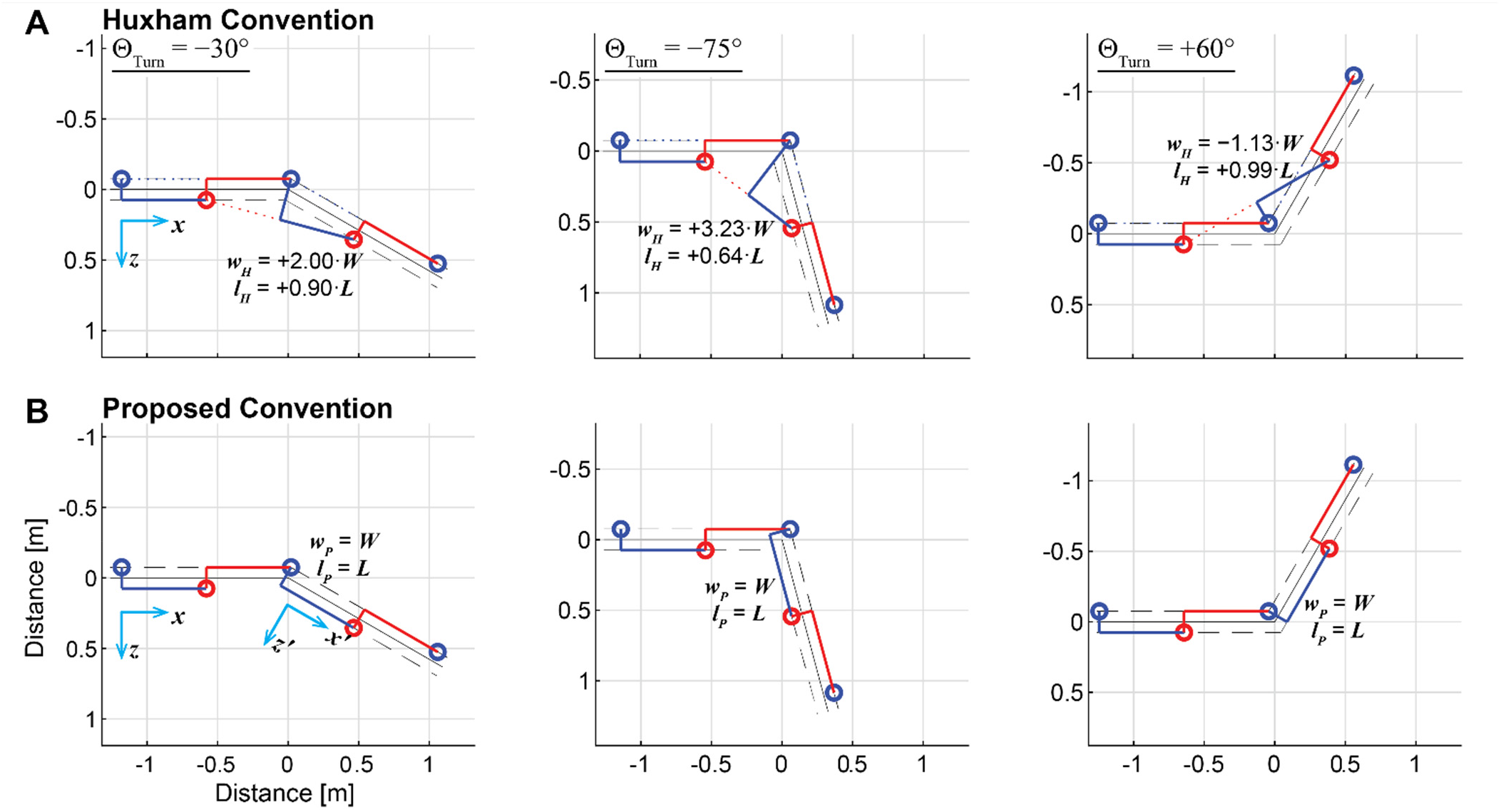
Single Discrete Turns: All example sequences shown were constructed to walk from left-to-right, executing a single turn at a single step at the 3^rd^ step shown. Top and bottom panels show the exact same sequences of *steps*. **A)** The Huxham defined “stride widths” (*w*_*H*_) and step lengths (*l*_*H*_) for stepping sequences involving turns executed at a single step. Huxham’s convention returns correct values (*w*_*H*_ = *W*; *l*_*H*_ = *L*) for all steps before and after the turn (not labeled). However, on the turn step itself, Huxham’s calculations return values that markedly deviate from these. Step width %Errors are+100%, +223%, and −213%, while step length %Errors are −10%, −36%, and −1%, for the 3 turns shown above. These errors scale with the amplitude of turn, despite the fact that the steps themselves (by construction) do not. **B)** The proposed step lengths (*l*_*P*_) and widths (*w*_*P*_) for the same stepping sequences. Here, because the local [*x*′,*z*′] coordinate system rotates with the turning movement on the step where the turn is executed, *all* steps in all sequences stepping return the correct values: i.e., *w*_*P*_ = *W* and *l*_*P*_ = *L* for all steps involved in all turns. The proposed methods thus yields 0 %Error for all measures for all steps, regardless of the magnitude of the turn executed.

For *Lateral Lane Change* maneuvers (Fig. 4), Huxham’s convention (Fig. 4A) yielded correct values (*l*_*H*_ = *L*; *w*_H_ = *W*) for only the first and last steps, but incorrectly distributes the maneuver (executed at a single step, by construction) across two consecutive steps (Fig. 4A), yielding counter-intuitive errors in both step lengths and widths at each step. Conversely, for our convention (Fig. 4B), because local path coordinates remain rightly aligned to lab coordinates (i.e., [*x′,z′*] = [*x,z*]) across all steps, correct step lengths and widths are obtained at every step (Fig. 4B).

**Figure 4.**
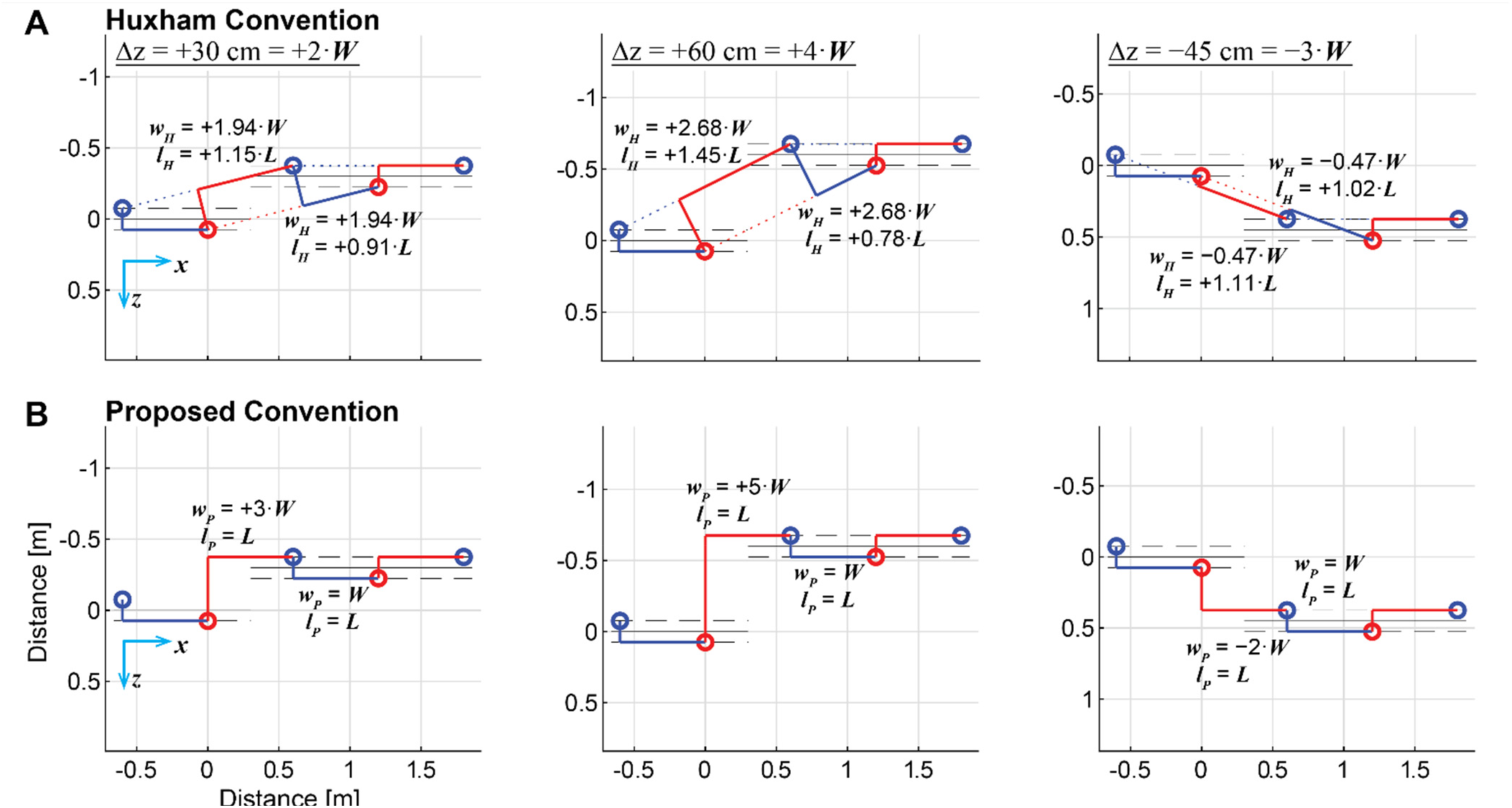
Lateral Lane-Change Maneuvers: All example sequences shown were constructed to walk from left-to-right, executing a single lateral lane change at a single step after the 2^nd^ step shown. Top and bottom panels show the exact same sequences of *steps*. **A)** The Huxham defined “stride widths” (*w*_*H*_) and step lengths (*l*_*H*_) for stepping sequences involving lateral lane-change maneuvers executed at a single step. Huxham’s convention returns correct values (*w*_*H*_ = *W*; *l*_*H*_ = *L*) for the first and last steps shown. However, despite that these stepping sequences were explicitly constructed to enact each lane change at a single step, Huxham’s convention necessitates distributing these maneuvers across two consecutive steps. In doing so, Huxham’s calculations return values that markedly deviate from defined movements that generated these lane changes. **B)** Conversely, the proposed convention keeps the local coordinate system ([*x*′,*z*′]) at each step aligned to the paths, which in turn change only location and not direction. Thus, all steps in all sequences yield exact step lengths of *l*_*P*_ = *L*. The proposed method likewise yields exact step widths of *w*_*P*_ = *W*+Δ*z* for the step that executes each lane change and *w*_*P*_ = *W* for all other steps in every sequence. The proposed methods thus yields 0 %Error for all measures for all steps, regardless of the magnitude of the lateral lane change executed.

For walking around *Circular Paths* (Figs. 5-6), both conventions yielded errors, as each approximated distances along a curve as straight lines. However, across all cases, Huxham’s convention produced far larger errors (Fig. 5A) than ours (Fig. 5B). These increased rapidly with decreasing *R*_*Path*_ (Fig. 6). Huxham’s convention yielded %Errors ∼20 *times* (range ∼2-40 times) larger for step lengths and ∼140 *times* (range ∼10-260 times) larger for step widths (Fig. 6A). These errors artificially introduced *large* stepping asymmetries (Fig. 6B) for steps purposefully constructed to be perfectly symmetrical. Conversely, our convention yielded exactly symmetrical (*USI* = 0) values (Fig. 6B), consistent with how stepping sequences were constructed.

**Figure 5.**
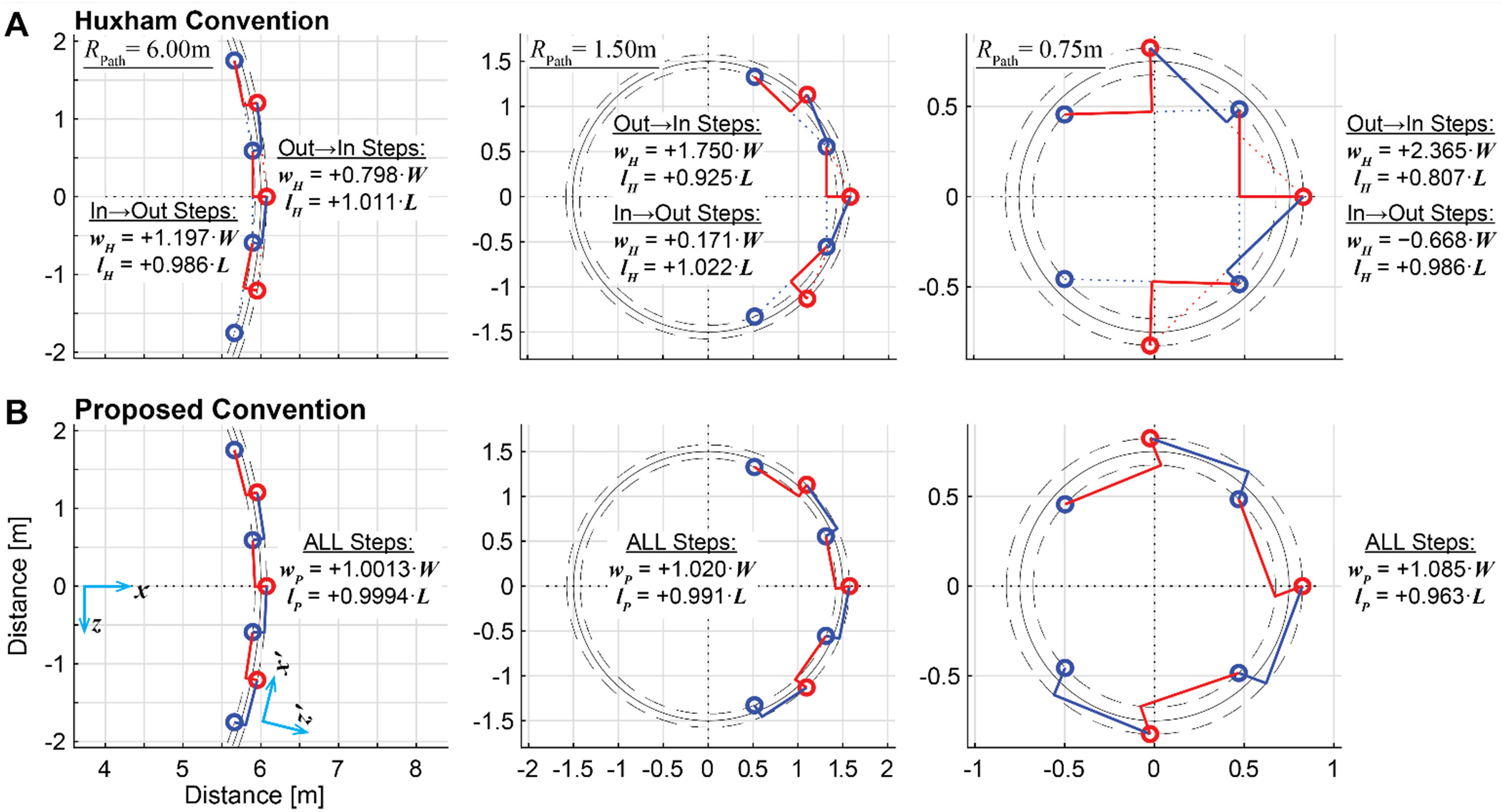
Continuous Walking Around a Circle: All example sequences shown were constructed to walk in a counter-clockwise direction around each circular path. Top and bottom panels show the exact same sequences of *steps*. **A)** The Huxham defined “stride widths” (*w*_*H*_) and step lengths (*l*_*H*_) for each stepping sequence. Because of how *w*_*H*_ and *l*_*H*_ are constructed (Fig. 1A), Huxham’s convention yields markedly different values for outside-to-inside (“out→in”; here, right-to-left) vs. inside-to-outside (“in→out”; here, left-to-right) steps. As shown, the deviations from the correct values (*w*_*H*_ = *W*; *l*_*H*_= *L*) increase dramatically as the radius of the circular path (*R*_*Path*_) gets smaller. **B)** The corresponding proposed step widths (*w*_*P*_) and lengths (*l*_*P*_) for the same stepping sequences. Because the proposed convention keeps the local coordinate system ([*x*′,*z*′]) at each step aligned tangent to the path at that step, and all steps were defined (by construction) equivalently, the proposed method yields the *same w*_*P*_ and *l*_*P*_ values for *all* steps, both outside-to-inside and inside-to-outside. While these values are not *exact* (due to approximating curvilinear motion as a straight line within each step), the deviations from the correct (defined) values (i.e., *w*_*P*_ = *W* and *l*_*P*_ = *L*) are clearly far smaller than for the Huxham convention.

**Figure 6.**
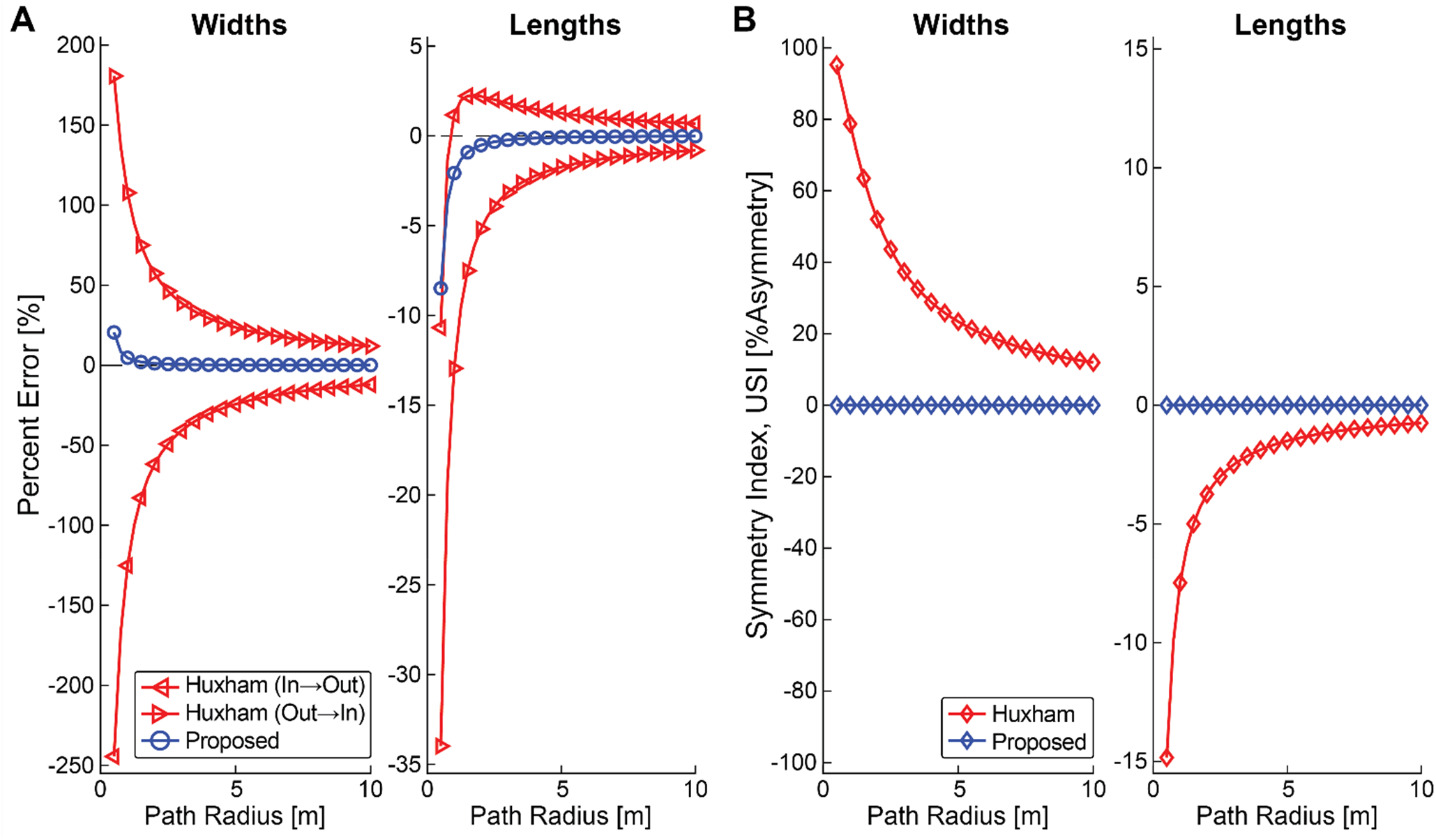
Stepping Percent Errors for Walking Around Circles: **A)** Percent stepping errors (%Error; Eq. (2), Methods) relative to their prescribed values, plotted versus circular path radius (*R*_*Path*_) for both step widths (left) and step lengths (right) for both the Huxham and proposed conventions. Note positive %Errors reflect larger-than-prescribed stepping parameter values, while negative %Errors reflect smaller-than-prescribed stepping parameter values. For very large path radii (i.e., as *R*_*Path*_ → ∞), both methods eventually converge to zero (0) %Error for straight walking. However, although %Errors for both methods tend to increase approximately exponentially as *R*_*Path*_ gets smaller, these errors are consistently at least ∼1-2 orders of magnitude smaller for our proposed convention (Fig. 1B) than for Huxham’s (Fig. 1A). **B)** Universal Symmetry Index (*USI*; Eq. (3), Methods) values reflecting relative asymmetries between inside-to-outside and outside-to-inside steps for the data shown in (A). Step widths and step lengths were both highly asymmetrical for the Huxham convention. Conversely, our proposed convention yielded perfectly symmetrical steps (consistent with how the steps were constructed), for all path radii.

Shifting [*x′,z′*] away from the mid-point between the feet (0.5; Fig. 1B), towards either the trailing foot (0.5 → 0; Fig. 2A) or leading foot (0.5 → 1; Fig. 2B), increased error magnitudes, with %Errors for outside-to-inside steps exhibiting trends opposite to those of inside-to-outside steps (Fig. 7A). This artificially introduced substantial asymmetries in both step lengths and widths (Fig. 7B) for steps purposefully constructed to be perfectly symmetrical (Fig. 5). As expected however, setting [*x′,z′*] at exactly 0.5 correctly yielded perfect symmetry (*USI*= 0; Fig. 7B), for any path curvature (Fig. 6B).

**Figure 7.**
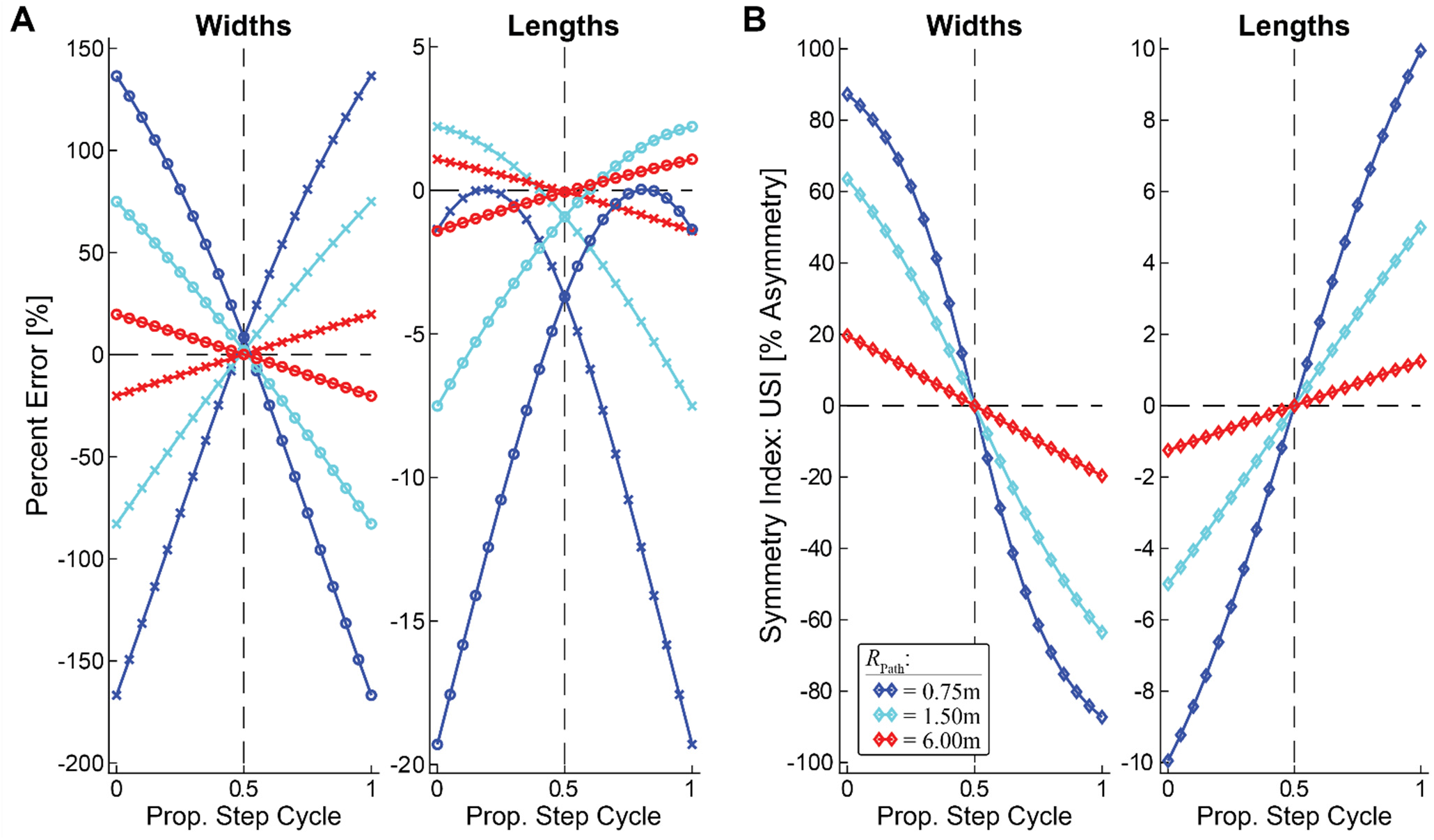
Stepping Percent Errors vs. [*x′,y′*] Location For Our Proposed Method: **A)** Percent stepping errors (%Error; Eq. (2), Methods) as computed with [*x′,y′*] set at different proportions of the gait cycle, starting from the location of the trailing foot (0; Fig. 2A) to the location of the leading foot (1; Fig. 2B). %Error data are for inside-to-outside (×) and outside-to-inside (?) steps for circular paths of the same path radii as in Fig. 4 (i.e. *R*_*Path*_ = {6.00, 1.50, 0.75} m). %Error magnitudes generally increased as [*x′,y′*] was shifted away from the mid-point between the feet (0.5) towards either the trailing foot (0.5→ 0) or leading foot (0.5 → 1). Trends for inside-to-outside steps were opposite to those of outside-to-inside steps. **B)** Universal Symmetry Index (*USI*; Eq. (3), Methods) values reflecting relative asymmetries between inside-to-outside and outside-to-inside steps for the data shown in **(A)**. Step widths and step lengths were symmetrical *only* when [*x′,y′*] was taken at the midpoint between the feet (0.5), and strongly *a*symmetrical otherwise, especially for step widths.

For walking along the *Arbitrary Curvilinear Path* (Fig. 8), both conventions yielded errors that generally scaled with the sharpness of local path curvature: errors were largest at/near the two sharpest turns towards the beginning and end of the path. However, across the entire path, our convention yielded errors (Fig. 8D) that were approximately order-of-magnitude smaller than Huxham’s (Fig. 8C).

**Figure 8.**
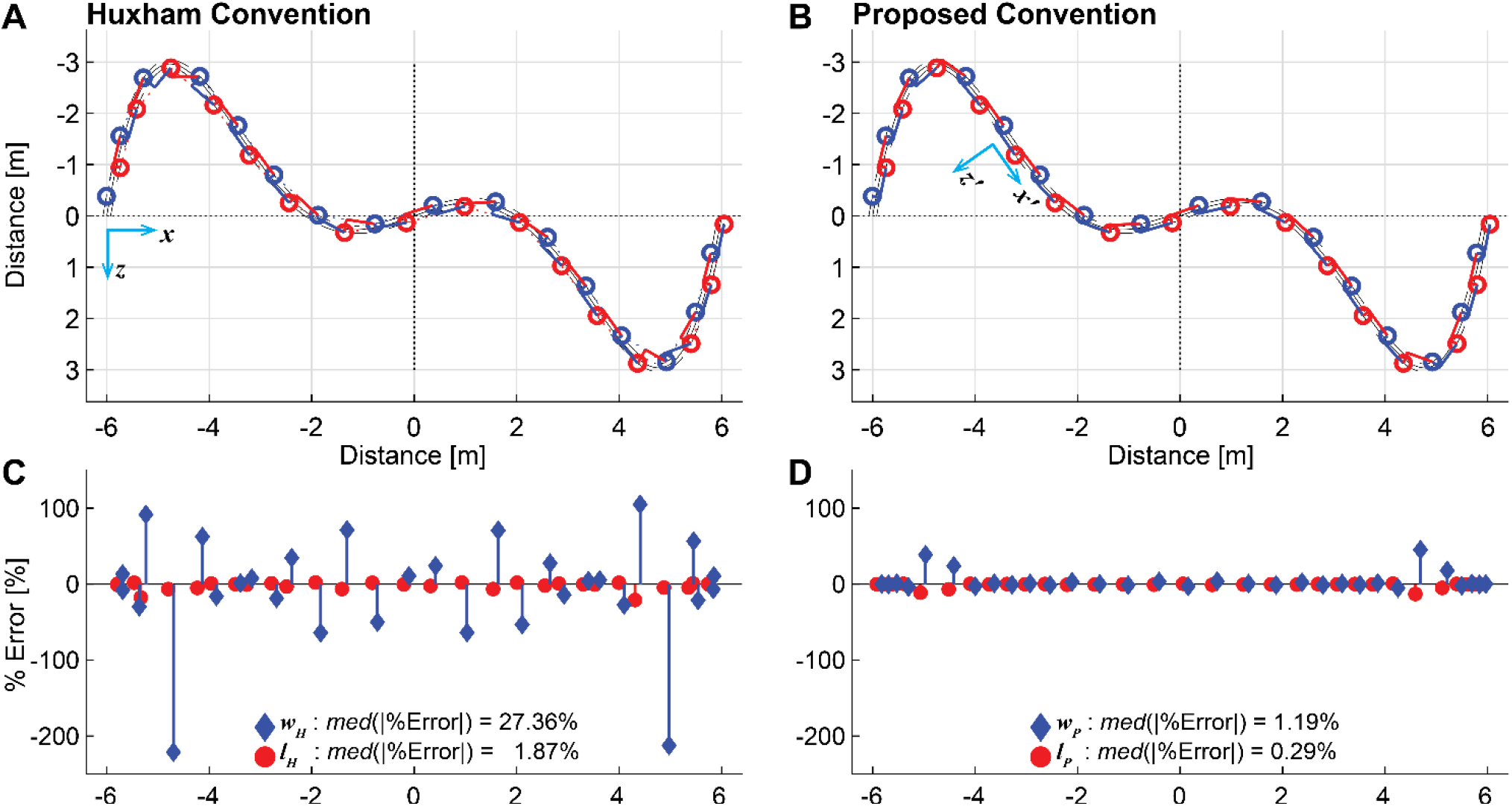
Continuous Walking Along an Arbitrary Path: A single sequence of 32 consecutive steps was constructed to walk along a curvilinear path defined as a 5^th^ order polynomial (see Methods). General direction of progression is from left to right. Panels (**A**) and (**B**) show the exact same sequences of *steps*. **A)** Huxham defined “stride widths” (*w*_*H*_) and step lengths (*l*_*H*_) for this stepping sequence. **B)** The corresponding proposed step widths (*w*_*P*_) and lengths (*l*_*P*_) for the same stepping sequence. **C)** %Errors at each step for both step widths (◊) and lengths (○), as computed following the Huxham convention.**D)** Corresponding %Errors at each step, as computed following the proposed convention. For both methods, the largest errors occurred at points of sharpest curvature, consistent with Figs. 4-5. Across all steps in this sequence, %Errors were consistently ∼order-of-magnitude smaller for the proposed convention than for the Huxham convention, also consistent with Figs. 4-5.. This was confirmed by computing the median (*med*) absolute %Error (i.e., |%Error|) across all steps in the sequence, as indicated on each plot.

## 4. Discussion

Real walking requires that we do more than just “take steps without falling” (Patil et al., 2022). Goal-directed walking is a *spatial navigation* task (Ekstrom et al., 2017). We walk intentionally (Gordon et al., 2021), to reach some destination or achieve other goals (Desmet et al., 2022). We navigate environments that are often complicated (Twardzik et al., 2019). This process is intrinsically anticipatory and requires considerable planning (Belmonti et al., 2013). As we walk, multiple cortical areas in our brains help estimate our location in our world and identify obstacles in our path (Drew and Marigold, 2015; Ekstrom et al., 2017). We use this information to plan our movements through our environment (Drew and Marigold, 2015; Wagner et al., 2014). These neural processes ultimately produce the walking paths we follow, whether relative to some explicitly-given path (store aisle, windy hiking trail, etc.), or the consistently stereotyped paths we construct for ourselves otherwise (Hicheur et al., 2007; Pham et al., 2011).

The work we present here underscores the theoretical necessity of knowing walking paths for measuring gait biomechanics. A significant conceptual advantage of our proposed convention is that it explicitly acknowledges one’s walking path to accurately estimate stepping parameters. Our proposed convention offers a coherent approach consistent with how humans plan and enact goal-directed walking neurophysiologically (Drew and Marigold, 2015; Ekstrom et al., 2017). Moreover, it is the simplest and most direct generalization of how biomechanists compute stepping parameters for straight walking, where we define steps relative to the task of walking along a straight-line path (Richards et al., 2023). Here, we still compute these foundational parameters in lab-based coordinates, only now we simply allow those lab-based coordinates to follow any non-straight path the person follows.

We show that anatomically-based coordinates cannot work because they intrinsically conflate body segment motions with the walking task performed (Supplement). The alternative, path-free convention proposed by (Huxham et al., 2006) (Fig. 1A) produces large artificial errors for single-step turns (Fig. 3A), introduces artificial changes in movement direction for single-step lateral maneuvers (Fig. 4A), generates both substantial errors (Fig. 5A & 6A) and artificial asymmetries (Fig. 6B) for walking around circles, and excessively large errors for walking on general curvilinear paths (Figs. 8A & 8C). Conversely, our convention (Fig. 1B) returns exactly correct stepping values for both turns (Fig. 3B) and lateral maneuvers (Fig. 4B), vastly smaller errors (Figs. 5B & 6A) and *no* artificial asymmetries (Figs. 6B & 7) for walking around circles, and vastly smaller errors for curvilinear walking (Figs. 8B & 8D).

One might consider a limitation to this study to be that we present no examples applied to experimental data. However, like prior efforts (Ho et al., 2023; Huxham et al., 2006; Kainz et al., 2016) that did so, we could only demonstrate that different conventions yield different values, because “true” values can never be known for experimental data. Thus, the strength of this work is precisely that we did use idealized simulated data to represent perfect performance for each walking task. Only by doing so could we quantify actual accuracy of each compared result relative to known *true* values. The essential conceptual questions our work answers could not have been addressed otherwise.

To this end, Figs. 3-4 simulate how a person could execute either a turn (Fig. 3) or lane-change maneuver (Fig. 4) with one step. While nothing biomechanical prevents this, real people typically take several steps to execute such turns (Conradsson et al., 2018; Dixon et al., 2013) or lane-changes (Acasio et al., 2017; Desmet et al., 2022). This does not invalidate what Figs. 3-4 demonstrate. Indeed, to simulate any such multi-step maneuver, we could simply concatenate together individual paths and corresponding steps, each of which executes some (perhaps small) turn (Fig. 3) or lateral shift (Fig. 4), the net sum of which yields the overall maneuver. Irrespective of the number of steps, if the (simulated) walker itself makes *no errors*, Figs. 3A & 4A demonstrate that any errors identified are attributable solely to Huxham’s convention.

One might also reason that another potential limitation to our convention (Fig. 1B) is needing to know the path the person follows. Numerous studies confirm that when people walk, they choose consistent (Moussaïd et al., 2011) and stereotypical paths (Hicheur et al., 2007; Huber et al., 2014; Pham and Hicheur, 2009). In lab experiments, walking tasks (and thus, paths) can be defined explicitly by the experimenter (Acasio et al., 2017; Bland et al., 2019; Desmet et al., 2022; Ho et al., 2023). Other lab-based studies may not impose explicit paths (Conradsson et al., 2018; Dixon et al., 2013; Madrid et al., 2023). When studying real-world walking (Bergsma et al., 2021; Glaister et al., 2007), defining explicit paths is less practical. As wearable measurement systems become more prevalent (Hafer et al., 2023), such technologies should seek to incorporate sensors to also situate the body *within its environment*, just as our brains do (Drew and Marigold, 2015).

One simple approach to identify walking paths post-collection is to filter out step-induced oscillations of body (e.g., center-of-mass, etc.) movements with a suitably low-cutoff (∼0.5 Hz) low-pass filter (Hicheur et al., 2007; Huber et al., 2014). Other applications may need to predict a person’s path *a priori* or in real time. Proposed approaches include physics-inspired “social force” models (Helbing et al., 2005; Helbing and Molnár, 1995), simple heuristic models that consider distance from, and time-to-collision to, upcoming obstacles (Moussaïd et al., 2011), or optimization approaches like maximizing path smoothness (Pham and Hicheur, 2009; Pham et al., 2007), minimizing variations in path curvature (Arechavaleta et al., 2008), or minimizing energetic cost (Brown et al., 2021). Many intelligent autonomous systems (like self-driving cars) also perceive and anticipate where humans will walk (Rudenko et al., 2020). Thus, numerous approaches exist to identify/estimate a person’s intended path. The convention we propose here offers a coherent approach to quantify stepping metrics relative to those paths.

Overall, our results strongly confirmed our hypothesis. Defining stepping parameters relative to the walking path (Fig. 1B) yielded *correct* results across more tasks, and more *accurate* results for each. Real world walking involves continuously making “embodied decisions” – we “decide as we act” while also “acting as we decide” (Gordon et al., 2021). To navigate complex spaces, people follow systematic, calculable trajectories (Arechavaleta et al., 2008; Moussaïd et al., 2011). They plan their walking paths (Drew and Marigold, 2015; Ekstrom et al., 2017), select foot placements several steps in advance (Matthis and Fajen, 2014) and regulate their steps from step to step to achieve specific task goals (Dingwell et al., 2010). One explicit goal is to *stay on their path* (Dingwell and Cusumano, 2019), which people readily modulate when their path changes (Desmet et al., 2022). Present findings are fully consistent with these principles and extend the argument for treating walking as a *goal-directed* process, where tasks are theoretically defined prior to, and independent of how people perform those tasks (Cusumano and Dingwell, 2013).

## Acknowledgments

This project was funded by the US National Institutes of Health / National Institute on Aging (NIA; Grant # R01-AG049735 & R21-AG053470; to JPC and JBD).

## References

Acasio, J., Wu, M.M., Fey, N.P., Gordon, K.E., 2017. Stability-maneuverability trade-offs during lateral steps. Gait Posture 52, 171–177.

Adolph, K.E., Cole, W.G., Komati, M., Garciaguirre, J.S., Badaly, D., Lingeman, J.M., Chan, G.L.Y., Sotsky, R.B., 2012. How Do You Learn to Walk? Thousands of Steps and Dozens of Falls per Day. Psychol. Sci. 23, 1387–1394.

Alves, S.A., Ehrig, R.M., Raffalt, P.C., Bender, A., Duda, G.N., Agres, A.N., 2020. Quantifying Asymmetry in Gait: The Weighted Universal Symmetry Index to Evaluate 3D Ground Reaction Forces. Frontiers in Bioengineering and Biotechnology 8.

Ambrose, A.F., Paul, G., Hausdorff, J.M., 2013. Risk factors for falls among older adults: A review of the literature. Maturitas 75, 51–61.

Arechavaleta, G., Laumond, J.P., Hicheur, H., Berthoz, A., 2008. An Optimality Principle Governing Human Walking. IEEE Trans Robot. 24, 5–14.

Belmonti, V., Cioni, G., Berthoz, A., 2013. Development of anticipatory orienting strategies and trajectory formation in goal-oriented locomotion. Exp. Brain Res. 227, 131–147.

Bergsma, B., Hulleman, D.N., Wiedemeijer, M.M., Otten, E., 2021. Foot placement variables of pedestrians in community setting during curve walking. Gait Posture 86, 120–124.

Bland, K., Lowry, K., Krajek, A., Woods, T., VanSwearingen, J., 2019. Spatiotemporal variability underlying skill in curved-path walking. Gait Posture 67, 137–141.

Brown, G.L., Seethapathi, N., Srinivasan, M., 2021. A unified energy-optimality criterion predicts human navigation paths and speeds. Proc. Natl. Acad. Sci. USA 118, e2020327118.

Bruijn, S.M., van Dieën, J.H., 2018. Control of human gait stability through foot placement. J. R. Soc. Interface 15, 1–11.

Burns, E.R., Kakara, R., 2018. Deaths from Falls Among Persons Aged ≥65 Years — United States, 2007–2016. Morb. Mortal. Wkly. Rep. 67, 509–514.

Conradsson, D., Paquette, C., Franzén, E., 2018. Medio-lateral stability during walking turns in older adults. PLoS ONE 13, e0198455.

Courtine, G., Schieppati, M., 2003. Human walking along a curved path. I. Body trajectory, segment orientation and the effect of vision. Eur. J. Neurosci. 18, 177–190.

Cusumano, J.P., Cesari, P., 2006. Body-Goal Variability Mapping in an Aiming Task. Biol. Cybern. 94, 367-379.

Cusumano, J.P., Dingwell, J.B., 2013. Movement Variability Near Goal Equivalent Manifolds: Fluctuations, Control, and Model-Based Analysis. Hum. Mov. Sci. 32, 899–923.

Desmet, D.M., Cusumano, J.P., Dingwell, J.B., 2022. Adaptive Multi-Objective Control Explains How Humans Make Lateral Maneuvers While Walking. PLoS Comput. Biol. 18, e1010035.

Dingwell, J.B., Cusumano, J.P., 2019. Humans Use Multi-Objective Control to Regulate Lateral Foot Placement When Walking. PLoS Comput. Biol. 15, e1006850.

Dingwell, J.B., John, J., Cusumano, J.P., 2010. Do Humans Optimally Exploit Redundancy to Control Step Variability in Walking? PLoS Comput. Biol. 6, e1000856.

Dixon, P.C., Stebbins, J., Theologis, T., Zavatsky, A.B., 2013. Spatio-temporal parameters and lower-limb kinematics of turning gait in typically developing children. Gait Posture 38, 870–875.

Drew, T., Marigold, D.S., 2015. Taking the next step: cortical contributions to the control of locomotion. Current Opinion in Neurobiology 33, 25–33.

Ekstrom, A.D., Huffman, D.J., Starrett, M., 2017. Interacting Networks of Brain Regions Underlie Human Spatial Navigation: A Review and Novel Synthesis of the Literature. J. Neurophysiol. 118, 3328–3344.

Fino, P.C., Nussbaum, M.A., Brolinson, P.G., 2016. Locomotor deficits in recently concussed athletes and matched controls during single and dual-task turning gait: preliminary results. J. Neuroeng. Rehabil. 13, 65.

Glaister, B.C., Bernatz, G.C., Klute, G.K., Orendurff, M.S., 2007. Video task analysis of turning during activities of daily living. Gait Posture 25, 289–294.

Gordon, J., Maselli, A., Lancia, G.L., Thiery, T., Cisek, P., Pezzulo, G., 2021. The road towards understanding embodied decisions. Neuroscience & Biobehavioral Reviews 131, 722–736.

Hafer, J.F., Vitali, R., Gurchiek, R., Curtze, C., Shull, P., Cain, S.M., 2023. Challenges and advances in the use of wearable sensors for lower extremity biomechanics. J. Biomech. 157, 111714.

Hase, K., Stein, R.B., 1999. Turning Strategies During Human Walking. J. Neurophysiol. 81, 2914–2922.

He, C., Xu, R., Zhao, M., Guo, Y., Jiang, S., He, F., Ming, D., 2018. Dynamic stability and spatiotemporal parameters during turning in healthy young adults. BioMedical Engineering OnLine 17, 127.

Helbing, D., Buzna, L., Johansson, A., Werner, T., 2005. Self-Organized Pedestrian Crowd Dynamics: Experiments, Simulations, and Design Solutions. Transportation Science 39, 1–24.

Helbing, D., Molnár, P., 1995. Social force model for pedestrian dynamics. Phys. Rev. E 51, 4282–4286.

Herssens, N., van Criekinge, T., Saeys, W., Truijen, S., Vereeck, L., van Rompaey, V., Hallemans, A., 2020. An investigation of the spatio-temporal parameters of gait and margins of stability throughout adulthood. J. R. Soc. Interface 17, 20200194.

Hicheur, H., Pham, Q.-C., Arechavaleta, G., Laumond, J.-P., Berthoz, A., 2007. The Formation of Trajectories During Goal-Oriented Locomotion in Humans. I. A Stereotyped Behaviour. Eur. J. Neurosci. 26, 2376–2390.

Ho, T.K., Kreter, N., Jensen, C.B., Fino, P.C., 2023. The choice of reference frame alters interpretations of turning gait and stability. J. Biomech. 151, 111544.

Huber, M., Su, Y.-H., Krüger, M., Faschian, K., Glasauer, S., Hermsdörfer, J., 2014. Adjustments of Speed and Path when Avoiding Collisions with Another Pedestrian. PLoS ONE 9, e89589.

Huxham, F., Gong, J., Baker, R., Morris, M., Iansek, R., 2006. Defining spatial parameters for non-linear walking. Gait Posture 23, 159–163.

Jansen, S.E.M., Toet, A., Werkhoven, P.J., 2011. Human locomotion through a multiple obstacle environment: strategy changes as a result of visual field limitation. Exp. Brain Res. 212, 449–456.

Kainz, H., Lloyd, D.G., Walsh, H.P.J., Carty, C.P., 2016. Instantaneous progression reference frame for calculating pelvis rotations: Reliable and anatomically-meaningful results independent of the direction of movement. Gait Posture 46, 30–34.

Kelsey, J.L., Procter-Gray, E., Hannan, M.T., Li, W., 2012. Heterogeneity of Falls Among Older Adults: Implications for Public Health Prevention. Am. J. Public Health 102, 2149–2156.

Lewis, C.L., Laudicina, N.M., Khuu, A., Loverro, K.L., 2017. The Human Pelvis: Variation in Structure and Function During Gait. The Anatomical Record 300, 633–642.

Madrid, J., Ulrich, B., Santos, A.N., Jolles, B.M., Favre, J., Benninger, D.H., 2023. Spatiotemporal parameters during turning gait maneuvers of different amplitudes in young and elderly healthy adults: A descriptive and comparative study. Gait Posture 99, 152–159.

Matthis, J.S., Fajen, B.R., 2014. Visual control of foot placement when walking over complex terrain. J. Exp. Psychol. Hum. Percep. Perf. 40, 106–115.

Moussaïd, M., Helbing, D., Theraulaz, G., 2011. How simple rules determine pedestrian behavior and crowd disasters. Proc. Natl. Acad. Sci. USA 108, 6884–6888.

Orendurff, M.S., Segal, A.D., Klute, G.K., Berge, J.S., Rohr, E.S., Kadel, N.J., 2004. The effect of walking speed on center of mass displacement. J. Rehabil. Res. Develop. 41, 829–834.

Patil, N.S., Dingwell, J.B., Cusumano, J.P., 2022. Viability, Task Switching, and Fall Avoidance of the Simplest Dynamic Walker. Scientific Reports 12, 8993.

Pham, Q.-C., Berthoz, A., Hicheur, H., 2011. Invariance of Locomotor Trajectories Across Visual and Gait Direction Conditions. Exp. Brain Res. 210, 207–215.

Pham, Q.-C., Hicheur, H., 2009. On the Open-Loop and Feedback Processes That Underlie the Formation of Trajectories During Visual and Nonvisual Locomotion in Humans. J. Neurophysiol. 102, 2800–2815.

Pham, Q.-C., Hicheur, H., Arechavaleta, G., Laumond, J.-P., Berthoz, A., 2007. The Formation of Trajectories During Goal-Oriented Locomotion in Humans. II. A Maximum Smoothness Model. Eur. J. Neurosci. 26, 2391–2403.

Prins, M.R., Cornelisse, L.E., Meijer, O.G., van der Wurff, P., Bruijn, S.M., van Dieën, J.H., 2019. Axial pelvis range of motion affects thorax-pelvis timing during gait. J. Biomech. 95, 109308.

Render, A.C., Kazanski, M.E., Cusumano, J.P., Dingwell, J.B., 2021. Walking humans trade off different task goals to regulate lateral stepping. J. Biomech. 119, 110314.

Richards, J., Levine, D., Whittle, M.W., 2023. Whittle’s Gait Analysis, 6th Ed. Elsevier.

Robinovitch, S.N., Feldman, F., Yang, Y., Schonnop, R., Leung, P.M., Sarraf, T., Sims-Gould, J., Loughin, M., 2013. Video capture of the circumstances of falls in elderly people residing in long-term care: an observational study. Lancet 381, 47–54.

Rudenko, A., Palmieri, L., Herman, M., Kitani, K.M., Gavrila, D.M., Arras, K.O., 2020. Human motion trajectory prediction: a survey. Int. J. Robotics Res. 39, 895–935.

Saunders, J.B.d.M., Inman, V.T., Eberhart, H.D., 1953. The Major Determinants in Normal and Pathological Gait. Journal of Bone and Joint Surgery 35-A, 543–558.

Taylor, M.E., Delbaere, K., Mikolaizak, A.S., Lord, S.R., Close, J.C.T., 2013. Gait parameter risk factors for falls under simple and dual task conditions in cognitively impaired older people. Gait Posture 37, 126–130.

Tesio, L., Rota, V., Chessa, C., Perucca, L., 2010. The 3D path of body centre of mass during adult human walking on force treadmill. J. Biomech. 43, 938–944.

Tillman, M., Molino, J., Zaferiou, A.M., 2022. Frontal plane balance during pre-planned and late-cued 90 degree turns while walking. J. Biomech. 141, 111206.

Twardzik, E., Duchowny, K., Gallagher, A., Alexander, N., Strasburg, D., Colabianchi, N., Clarke, P., 2019. What features of the built environment matter most for mobility? Using wearable sensors to capture real-time outdoor environment demand on gait performance. Gait Posture 68, 437–442.

Wagner, J., Solis-Escalante, T., Scherer, R., Neuper, C., Müller-Putz, G., 2014. It’s how you get there: walking down a virtual alley activates premotor and parietal areas. Frontiers in Human Neuroscience 8, 93.

Xu, R., Wang, X., Yang, J., He, F., Zhao, X., Qi, H., Zhou, P., Ming, D., 2017. Comparison of the COM-FCP inclination angle and other mediolateral stability indicators for turning. BioMedical Engineering OnLine 16, 37.

