## Supplement for "Generalizing Stepping Concepts To Non-Straight Walking"

### SUPPLEMENT — ADDITIONAL RESULTS

*Why Using "Anatomically-Based" Coordinates Is Not Viable*

Some have suggested using anatomically-based local coordinates systems, including those defined by the lateral CoM trajectory (Ho et al., 2023) or transverse (i.e., horizontal) plane pelvis (Christensen et al., 2023; Ho et al., 2023; Kainz et al., 2016) or trunk (Jansen et al., 2011) angles to identify the direction of forward progression ("DoP") of the body during each step. Here, we show why these approaches are not viable for computing *task*-relevant variables like stepping parameters (i.e., step lengths and widths), even for the supposedly "simplest" case of walking in a straight line.

The confounds arise from basic gait mechanics long-known since the foundational work of Saunders (Saunders et al., 1953). Even when walking in a straight line, the pelvis (and CoM) oscillates side-to-side as humans shift their weight over each new stance foot (Saunders' $6^{th}$ "determinant of gait"). The pelvis also rotates in the transverse plane (Lewis et al., 2017) (Saunders' $1^{st}$ "determinant of gait"), helping to extend step length (Liang et al., 2014). The trunk (thorax) likewise rotates horizontally relative to the pelvis (Lamoth et al., 2002; Prins et al., 2019b), etc. Human walking involves complex rotations that interact across multiple segments. Thus, no individual segment motion can uniquely define "the body" (as a whole).

Here, we simulated the task of taking "perfect" steps (all $L$ = 0.60m; $W$ = 0.15m) to walk in a straight line (i.e., in the direction of the $+x$-axis; Fig. S1A). We first approximated the lateral center-of-mass (CoM) motion trajectory (Desmet et al., 2022; Saunders et al., 1953; Tesio et al., 2010; Winter, 1995) as a cosine function (Fig. S1A):

$$z_{CoM} = \left(0.8\frac{W}{2}\right)\cdot\cos\left(\frac{180°}{L}\cdot x\right). \tag{1}$$

We computed the corresponding horizontal plane angle of orientation of the tangent to this curve to define the instantaneous direction of motion of the pelvis (Fig. S1B).

We then approximated the transverse plane rotation of the pelvis (Lewis et al., 2017; Prins et al., 2019b) as a double-cosine function (Fig. S1B):

$$\theta_{Pelvis} = -4.5\cdot\cos\left(\frac{180°}{L}\cdot(x-0.2)\right)+1.5\cdot\cos\left(3\cdot\frac{180°}{L}\cdot(x-0.2)\right). \tag{2}$$

We then also approximated the transverse plane rotation of the thorax (Lamoth et al., 2002; Prins et al., 2019b) also as a (somewhat different) double-cosine function (Fig. S1B):

$$\theta_{Thorax} = -2.5\cdot\cos\left(\frac{180°}{L}\cdot(x-0.2)\right)-0.5\cdot\cos\left(3\cdot\frac{180°}{L}\cdot(x-0.2)\right). \tag{3}$$

Though approximate, these curves are representative. Their precise amplitudes, shapes, and relative timing is not essential to the central question(s) addressed here. The question addressed here is to what extent can any such curve be used to define a person's DoP on a given step, which is necessary as precursor to defining step widths and/or lengths.

For walking humans, the amplitudes and shapes of the above trajectories vary with multiple factors, including (among others) sex (Lewis et al., 2017), gait speed (Lamoth et al., 2002; Liang et al., 2014; Prins et al., 2019b; Tesio et al., 2010), arm swing (Prins et al., 2019b), trunk stiffness (Prins et al., 2019a; Prins et al., 2019b), pathology (Lamoth et al., 2002; Lewis et al., 2017), etc. Thus, any directions specified from these curves will align with the *true* DoP at different points in the step cycle that will themselves vary accordingly. When these alignments will occur is not knowable *a priori*.

Here, we used simulated steps and anatomical rotation curves to directly quantify actual *accuracy* of calculations. The accuracy of any measure can only be computed relative to *known true values* (here, $L = 0.60$m; $W = 0.15$m). Such analyses cannot be conducted with experimental data, as true values cannot be known. To our knowledge, this work is therefore the first to directly quantify the actual (potential/likely) error magnitudes associated with using anatomically-based coordinate reference frames [as in e.g., (Ho et al., 2023; Jansen et al., 2011; Kainz et al., 2016)] to estimate step lengths and widths.

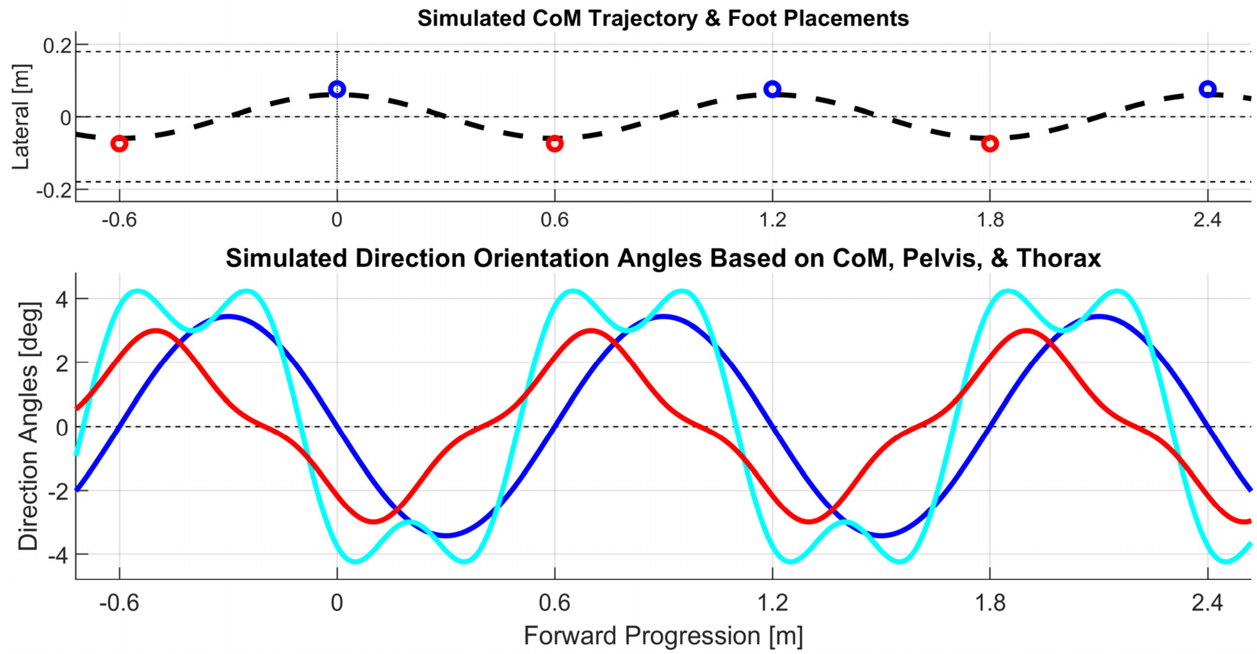


**Figure S1: Angular Directions From Local Coordinate Choices. A:** Left (**O**) and Right (**O**) foot placements for taking perfect steps walking in a straight line. Dashed line simulates approximate CoM trajectory (Eq. 1). **B:** Simulated transverse plane direction angles for approximate CoM Motion (▬; Eq. 1), pelvis angles (▬; Eq. 2), and thorax angles (▬; Eq. 3).

Here, we quantified these effects by varying the location of the local coordinates [$x'$, $y'$] to be aligned to the angles of each curve (Fig. S1B) proportionally (i.e., from 0 to 1) from the location of the trailing foot (0) to the location of the leading foot (1) of each step. At each [$x'$, $y'$] location, we computed step widths and step lengths in [$x'$, $y'$] coordinates (Fig. S2) and %Errors relative to known true values in each (Fig. S3).

Both step widths and step lengths deviated substantially from known true values ($L = 0.60$m; $W = 0.15$m) at nearly every point in the step cycle (Fig. S2). These corresponded to %Errors generally in the range of ± ~1-2% for step lengths, but upwards of ± ~30% for step widths (Fig. S3). Each local coordinate choice (CoM vs. pelvis vs. thorax) perfectly aligned with the true DoP (i.e., $+x$ direction), and thus exhibited zero (0) %Error, generally only once during each step cycle and (importantly) at different relative points in time during the step cycle that were not easily *a priori* discernable.

To use any of these candidate anatomical coordinate frames, one must select some instant during the step cycle to set the local coordinates, [$x'$, $y'$]. Figs. S2 & S3 make it abundantly apparent that step length / width calculations made in any such frame will depend greatly on the phase of the step cycle chosen. One can only obtain the correct answers when the candidate variable is known to be directly aligned in the same direction as the true DoP. One can only know this if one already knows (or can closely estimate, as in Arechavaleta et al. (2008), Moussaïd et al. (2011), etc.) the (*task*-defined) walking trajectory or *path*, independent of the motions of the individual body segments. However, having to know this path *a priori* effectively negates the intended purpose of adopting any of these above-described anatomically-based alternatives to begin with.

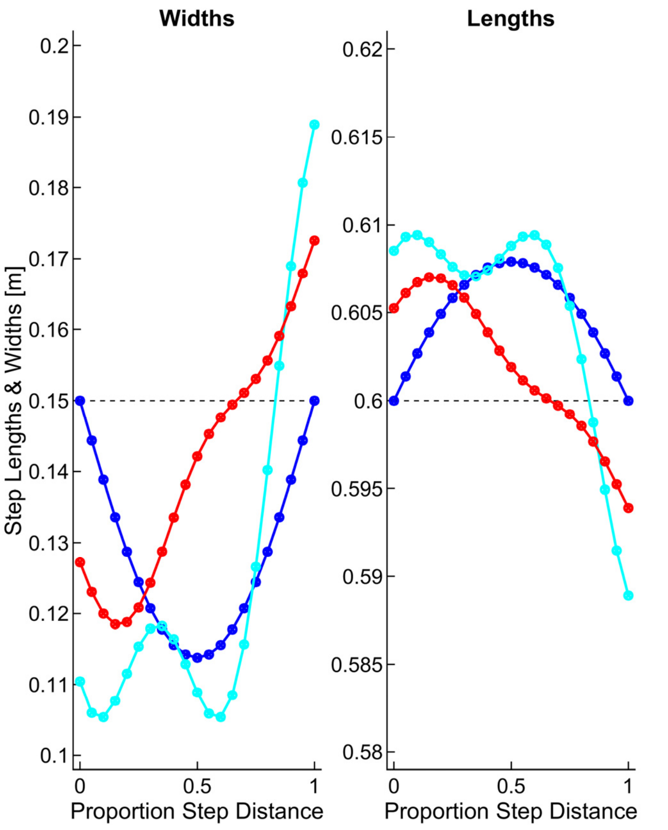


**Figure S2: Step Lengths & Widths vs. [*x′*, *y′*] Location.** Data shown are for [*x′*, *y′*] defined by CoM Motion (▬; Eq. 1), pelvis angles (▬; Eq. 2), and thorax angles (▬; Eq. 3).

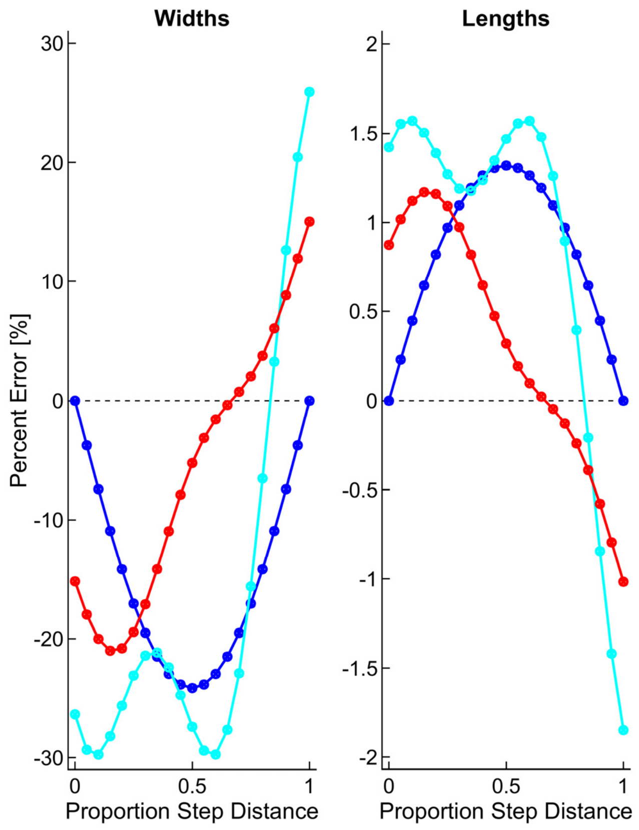


**Figure S3: Percent Errors vs. [*x′*, *y′*] Location.** Data shown are for the step widths and lengths shown in Fig. S2-2.

The same types of results are observed experimentally: using differently-defined coordinate reference frames and/or setting coordinates at different phases of the step cycle yields different results for *task*-relevant variables, including step widths and lengths and also Margins of Stability (MoS) (Ho et al., 2023). The differences reflected in such results are essentially arbitrary and cannot be reconciled if/when the *true* DoP is not known. This is because these anatomical motions of the CoM, pelvis, trunk (or other segment), reflect *how* people move as they perform the task of walking. However, they do *not* reflect the task itself that these people are performing. The task to be performed (here, walk in a straight line) is defined external to, and independent of, the movements we enact to accomplish that task (Cusumano and Cesari, 2006).

This makes all the preceding anatomically-based coordinate system options highly problematic, not only for non-straight walking (Christensen et al., 2023; Ho et al., 2023; Kainz et al., 2016), but even for simple straight walking. These results obtained are clearly arbitrary (Figs. S2-S3). Thus, it cannot be recommended to adopt any of the approaches mentioned above, or any other similar such anatomically-defined coordinate system to assess stepping.

### References:


Arechavaleta, G., Laumond, J.P., Hicheur, H., Berthoz, A., 2008. An Optimality Principle Governing Human Walking. IEEE Trans Robot. 24, 5-14.

Christensen, M.S., Tracy, J.B., Crenshaw, J.R., 2023. A pelvic-oriented margin of stability is robust against deviations in walking direction. SportRχiv [Pre-Print].

Cusumano, J.P., Cesari, P., 2006. Body-Goal Variability Mapping in an Aiming Task. Biol. Cybern. 94, 367-379.

Desmet, D.M., Cusumano, J.P., Dingwell, J.B., 2022. Adaptive Multi-Objective Control Explains How Humans Make Lateral Maneuvers While Walking. PLoS Comput. Biol. 18, e1010035.

Ho, T.K., Kreter, N., Jensen, C.B., Fino, P.C., 2023. The choice of reference frame alters interpretations of turning gait and stability. J. Biomech. 151, 111544.

Jansen, S.E.M., Toet, A., Werkhoven, P.J., 2011. Human locomotion through a multiple obstacle environment: strategy changes as a result of visual field limitation. Exp. Brain Res. 212, 449-456.

Kainz, H., Lloyd, D.G., Walsh, H.P.J., Carty, C.P., 2016. Instantaneous progression reference frame for calculating pelvis rotations: Reliable and anatomically-meaningful results independent of the direction of movement. Gait Posture 46, 30-34.

Lamoth, C.J.C., Meijer, O.G., Wuisman, P.I.J.M., van Dieën, J.H., Levin, M.F., Beek, P.J., 2002. Pelvis-Thorax Coordination in the Transverse Plane During Walking in Persons With Nonspecific Low Back Pain. Spine 27.

Lewis, C.L., Laudicina, N.M., Khuu, A., Loverro, K.L., 2017. The Human Pelvis: Variation in Structure and Function During Gait. The Anatomical Record 300, 633-642.

Liang, B.W., Wu, W.H., Meijer, O.G., Lin, J.H., Lv, G.R., Lin, X.C., Prins, M.R., Hu, H., van Dieën, J.H., Bruijn, S.M., 2014. Pelvic step: The contribution of horizontal pelvis rotation to step length in young healthy adults walking on a treadmill. Gait Posture 39, 105-110.

Moussaïd, M., Helbing, D., Theraulaz, G., 2011. How simple rules determine pedestrian behavior and crowd disasters. Proc. Natl. Acad. Sci. USA 108, 6884-6888.

Prins, M.R., Bruijn, S.M., Meijer, O.G., van der Wurff, P., van Dieën, J.H., 2019a. Axial Thorax-Pelvis Coordination During Gait is not Predictive of Apparent Trunk Stiffness. Scientific Reports 9, 1066.

Prins, M.R., Cornelisse, L.E., Meijer, O.G., van der Wurff, P., Bruijn, S.M., van Dieën, J.H., 2019b. Axial pelvis range of motion affects thorax-pelvis timing during gait. J. Biomech. 95, 109308.

Saunders, J.B.d.M., Inman, V.T., Eberhart, H.D., 1953. The Major Determinants in Normal and Pathological Gait. Journal of Bone and Joint Surgery 35-A, 543-558.

Tesio, L., Rota, V., Chessa, C., Perucca, L., 2010. The 3D path of body centre of mass during adult human walking on force treadmill. J. Biomech. 43, 938-944.

Winter, D.A., 1995. Human Balance And Posture Control During Standing And Walking. Gait Posture 3, 193-214.
